# G Protein Coupled Estrogen Receptor Signaling Maintains β Cell Identity in Female Mice

**DOI:** 10.1101/2025.05.12.652914

**Authors:** Madeline R. McLaughlin, Preethi Krishnan, Wenting Wu, Cameron Rostron, Kara Orr, Lata Udari, Jacqueline Del Carmen Aquino, Amanda Fisher, Tatsuyoshi Kono, Kok Lim Kua, Carmella Evans-Molina

## Abstract

Type 2 diabetes (T2D) arises in the context of obesity and overnutrition; however, additional demographic features including age and biological sex contribute to T2D risk. Estradiol (E2) is thought to play a protective metabolic role that may govern sex differences in the development of T2D. The mechanisms by which E2 exerts these effects and the impact of reduced E2 signaling in β cells during menopause remain incompletely understood. We analyzed publicly available whole islet transcriptome datasets from female and male cadaveric donors and showed significant age-related modulation of gene expression, including changes in pathways related to β cell function, in islets from female donors. Importantly, these patterns were not observed in islets from male donors. To test the *in vivo* relationship between E2 signaling and β cell function, 10-week-old female C57BL6/J mice underwent an ovariectomy (OVX) or sham (CTR) surgery, followed by 4 weeks of high-fat diet (HFD) treatment. HFD-OVX mice exhibited obesity-induced glucose intolerance, increased α cell mass, and reduced expression of β cell identity markers. Furthermore, *ex vivo* treatment of islets with the G protein coupled estrogen receptor (GPER)- specific agonist G-1 restored β cell identity gene expression. Together, these data identify a novel connection between GPER signaling and β cell identity and suggest that menopausal loss of E2 signaling through GPER may be linked with loss of β cell identity.

## 1. Introduction

Type 2 diabetes (T2D) mellitus is a global health pandemic that accounts for 90% of all forms of diabetes and affects more than 537 million individuals worldwide (1). The pathophysiology of T2D is characterized by a combination of peripheral insulin resistance and pancreatic β cell dysfunction, while features such as obesity, overnutrition, lack of physical activity, inflammation, aging, and genetics modify this pathophysiology and contribute to an individual’s overall risk of developing T2D (2). In addition to these genetic, demographic, and environmental factors, biological sex also governs the development of T2D across the lifespan. Across multiple ethnic groups, men have an increased risk for developing T2D as compared to pre-menopausal women (3–6). Worldwide, 17.7 million more men than women have been diagnosed with T2D. Moreover, men are diagnosed at a younger age and have lower body fat mass than women on average (7).

However, in midlife, women have an increased risk of developing T2D compared to younger women, which is thought to occur secondary to menopause. The average age of menopause in women in the United States is 51 years of age, and menopause marks the transition when a woman’s ovaries no longer produce endogenous estrogen (E2). Due to this lack of E2 signaling, a variety of physiological changes are observed, including alterations in body composition (i.e., increased visceral adiposity), reduced insulin sensitivity, inadequate insulin secretion, and increased inflammation (8, 9). Post-menopausal women and women who have undergone surgical menopause (i.e., bilateral oophorectomy) exhibit an increased risk for T2D that is comparable to the risk seen for men. Additionally, treatment of post-menopausal women with E2 replacement therapy is associated with reduced diabetes risk, highlighting the importance of both aging and E2 signaling in metabolism and the development of T2D (10).

E2 is an important regulator of whole-body physiology and is known to exert protective metabolic effects through its action at the three E2 receptors: estrogen receptor α (ERα), estrogen receptor β (ERβ), and G protein-coupled estrogen receptor 1 (GPER1/GPR30). ERα and ERβ are the classical nuclear and extranuclear estrogen receptors that mediate transcriptional and genomic activities and activate a variety of signaling cascades to facilitate nongenomic effects (11). ERα and ERβ are the most well-studied estrogen receptors and serve as essential regulators of glucose and lipid metabolism. Activation of ERα has been shown to enhance glucose-stimulated insulin biosynthesis, reduce de novo synthesis of fatty acids and lipogenesis, prevent accumulation of toxic lipid intermediates, and promote β cell survival in response to proapoptotic stimuli associated with diabetes (12–18). Global ERα deletion in female mice results in elevated fasting insulin levels, impaired glucose tolerance, and increased skeletal muscle insulin resistance (19, 20).

GPER, though less well-studied, is also an essential mediator of nongenomic signaling through E2 as evidenced by the fact that E2 is still able to exhibit anti-diabetic actions in mice with deletion of both ERa and ERb (18). Notably, total body GPER knockout (KO) in female mice leads to glucose intolerance and impaired insulin secretion (19–23). Further, in a female ovariectomy mouse model of post-menopausal obesity, treatment with the GPER specific agonist G-1 reduced body weight, improved glucose homeostasis, increased energy expenditure, and decreased body fat content with a subsequent reduction in fasting cholesterol, glucose, insulin, and inflammatory markers, suggesting that GPER-selective agonism is a potential therapeutic target for treating metabolic disorders such as obesity and diabetes (24).

We demonstrated recently that genetic loss of STIM1 leads to loss of GPER expression and reduced β cell function and identity in female mice (25), raising the question of whether surgical or physiological cessation of E2 signaling is similarly associated with changes in β cell identify. To test this hypothesis, we evaluated islet RNA sequencing datasets from male and female organ donors, and we performed ovariectomy (OVX) surgery coupled with short-term high fat diet (HFD) exposure in female mice.

## 2. Methods

### 2.1 Human Islet RNA-Sequencing Analysis

To test the impact of age on human islet gene expression in male and female organ donors, GSE152111 was retrieved from the NCBI Gene Expression Omnibus database. The dataset included RNA sequencing data from 66 human pancreatic islet donors. RNA sequencing datasets were generated using Illumina HiSeq 2500 sequencing. Male (n=40) and female (n=26) samples were analyzed separately. Pearson correlation was performed on genes that had a normalized value of at least 1 on average. Genes that showed r>0.4 and p<0.05 were considered to be correlated with age. The dataset was further filtered to retain only the protein-coding genes. The correlated genes were compared between males and females, and functional enrichment analysis was performed separately for male-specific, female-specific, and common genes using the DAVID bioinformatics tool (26, 27). Specifically, biological processes from gene ontology analyses were identified. Terms with fold enrichment value ≥1.3 and p<0.05 were considered.

### 2.2 Animals

Female C57Bl/6J wildtype (WT) mice were purchased from Jackson Laboratory at 7 weeks of age and allowed to acclimate for 3 weeks. To model E2 deficiency, bilateral ovariectomy (OVX) or sham (CTR) surgeries were performed in our facility at 10 weeks of age (28). These studies were approved by the Indiana University School of Medicine Animal Care and Use Committee (#24134). Sham CTR mice were treated identically, apart from OVX. To validate that the OVX surgery was effective, serum E2 levels were measured 5 weeks post-surgery using liquid chromatography-mass spectroscopy (lower limit of detection: 0.00125 ng/mL (29)) at the Wisconsin National Primate Research Center Assay Services Unit (University of Wisconsin).

### 2.3 Metabolic Studies

At 9 weeks of age (1 week prior to surgery), mouse body weight and composition were determined using the EchoMRI-500 Body Composition Analyzer (EchoMRI; RRID:SCR_017104) to obtain baseline measurements. Glucose tolerance tests (GTT) were performed after a 6-hour fast followed by intraperitoneal (IP) administration of 1.5 g D-glucose/kg lean mass. Insulin tolerance tests (ITT) were performed after 6 hours of fasting followed by IP administration of 0.75 U Novolin insulin per kg lean mass (Novo Nordisk). Blood glucose levels were measured using AlphaTrak® 3 disposable test strips (10024743, ADW Diabetes) and an AlphaTrak® 3 glucometer (10024742), and OVX or CTR surgeries were performed at 10 weeks of age. After surgery, mice were injected with 20 mg/kg carprofen (RIM-00284R1, Zoetis) once daily over three consecutive days for analgesia. Directly following the surgery, mice were maintained on a normal chow diet (2019SX, Inotiv) for 1 week while they recovered. Beginning at 1-week post-surgery (11 weeks of age), mice were fed *ad libitum* with a HFD (45% kcal from fat, Research Diets, catalog #D12451). After 4 weeks of HFD, body weight and body composition were again analyzed using EchoMRI, and GTTs and ITTs were performed, as described above.

### 2.4 Isolation of Murine Islets

Pancreatic islets of experimental animals were isolated by the Indiana University School of Medicine Islet & Physiology Core after 5 weeks of HFD (6 weeks post-surgery) by collagenase digestion, as previously described (30). Briefly, following injection of type XI collagenase (C7657, Sigma-Aldrich) into the common bile duct, the pancreas was excised and digested. Islets were retrieved using a Histopaque (10771, 11191, Sigma-Aldrich) gradient (30). Islets were dispersed in phenol red-free, 5.5 mM low-glucose DMEM (11054-020, Gibco) supplemented with 10% charcoal-stripped fetal bovine serum (35-072-CV, Corning), 10 mM HEPES (15630130, Gibco), 100 U/mL penicillin-streptomycin (15140122, Gibco), and 2 mM L-glutamine (25030081, Gibco). Islets were then handpicked and cultured in the media listed above at 37 °C with 5% CO_2_ for overnight recovery. All *ex vivo* islet assays were performed following an overnight recovery period unless stated otherwise.

### 2.5 Quantitative RT-PCR

Approximately 50-75 isolated mouse islets were washed with PBS, and total RNA was extracted using the RNeasy Micro Plus Kit (74034, Qiagen) per the manufacturer’s recommendations. The entire elute (∼11 μL) was reverse transcribed using Moloney Murine Leukemia Virus Reverse Transcriptase (28025013, Invitrogen) following manufacturer’s recommendations. cDNA was subjected to RT-qPCR using SensiFAST SYBR Lo-ROX (BIO- 94050, Bioline) and a QuantStudio 3 thermocycler (A28567, Applied Biosystems). PCR reactions were heated to 98 °C for 5 minutes to denature the DNA and activate the polymerase. Reactions were then cycled 40 times from 98 °C for 15 sec to 60 °C for 1 minute, then increasing temperature at 0.15 °C/sec with signal acquisition until 95 °C was reached. Relative RNA levels were established against *Actb* mRNA using the comparative ΔΔCt method, as previously described (31). Primer sequences are provided in Table 1.

**Table 1:**
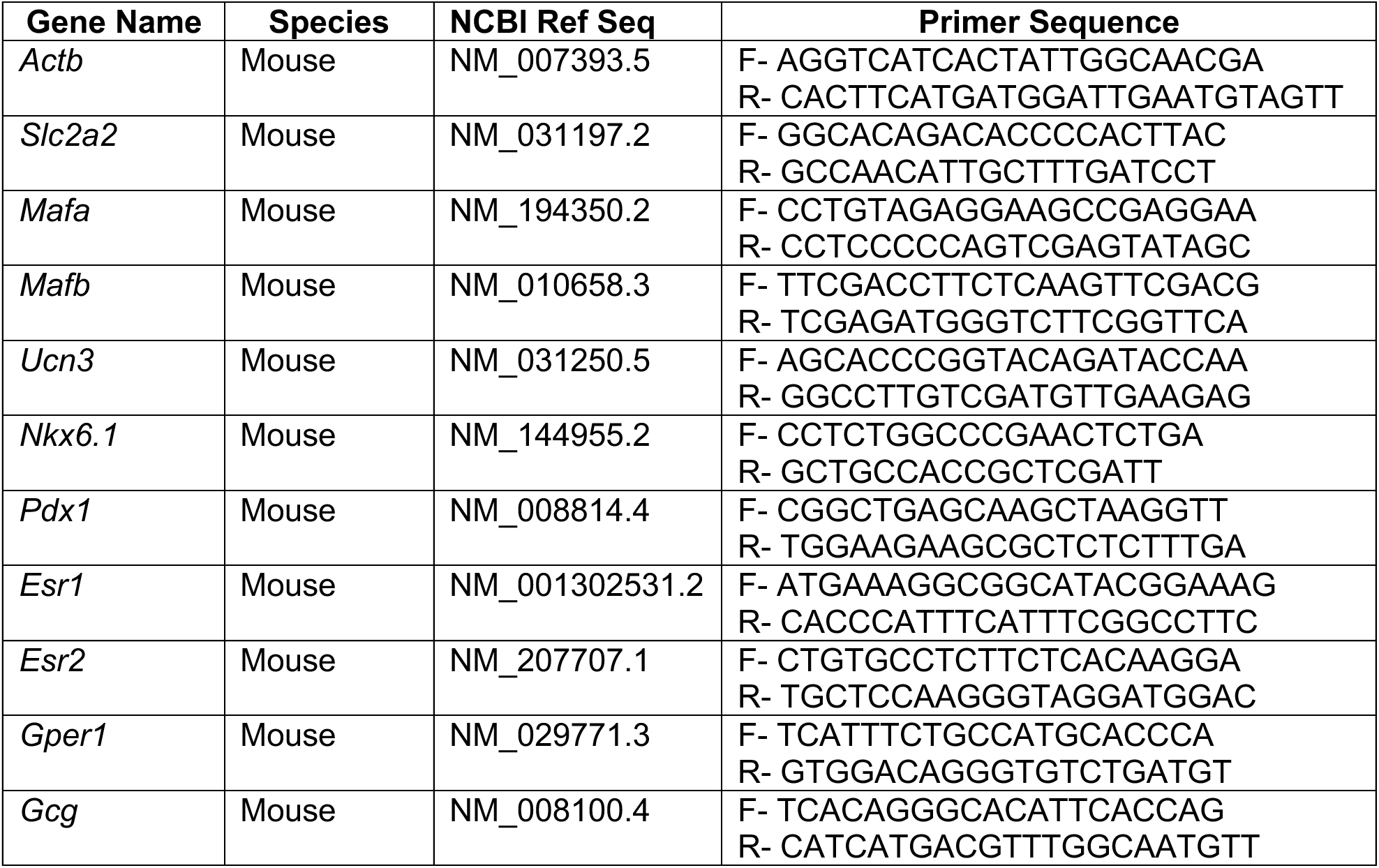
Primer List for RT-qPCR.

### 2.6 Treatment with E2 and GPER1 Selective Agonist G-1

Islets isolated from female mice that underwent OVX or CTR procedure were plated in non-treated 6-well plates at a concentration of 75 islets per well. Twenty-four hours after isolation, medium was replaced with fresh charcoal-stripped medium or charcoal-stripped medium containing 0.1 μM E2 (E0950, Steraloids) or 0.1 μM GPER1 selective agonist G-1 (10008933, Caymen Chemicals) (25, 32). Islet cells were cultured for 24 hours and collected for total RNA isolation and RT-qPCR analysis, as described above.

### 2.7 Immunofluorescence and Integrative Density Analysis of Mouse Pancreas

After euthanasia, mouse pancreata were rapidly removed and fixed overnight in Z-fix buffered zinc formalin fixative (Anatech Ltd.). Fixed specimens were paraffin-embedded and 5 μm thick longitudinal sections were collected. Following deparaffinization, rehydration, and antigen retrieval by heating in a citrate antigen retrieval buffer (H-3300-250, Vector Laboratories), sections were permeabilized, blocked with Animal-Free Blocker (SP-5030-250, Vector Laboratories), and incubated with primary antibodies overnight at 4 °C (Insulin – Dako, A0564, RRID:AB_10013624, Glucagon – Abcam, ab92517, RRID:AB_10561971, Ki67 – Cell Signaling Technology, #9129, RRID:AB_2687446). Incubation with secondary antibodies conjugated to AlexaFluor (Thermo Fisher) was performed at room temperature for 1.5 hours. Sections were mounted with Fluorosave (345789, Millipore). Z-stack images of islets were acquired using a Zeiss confocal microscope LSM 800 (Carl Zeiss, RRID:SCR_015963) using a 40X oil immersion objective. Images were analyzed with ImageJ, Fiji software (NIH, RRID:SCR_002285). Fluorescence intensity of insulin and glucagon (number of pixels) was calculated by measuring integrated density (ID) and total islet area. The background fluorescence intensity was measured for each channel, and the corrected total islet cell fluorescence intensity was calculated by the following formula: total islet cell fluorescence (CTCF) = integrated density – (area of selected cell x mean fluorescence of background) (33).

The mass of β and α cells was estimated for each animal by determining the average β cell and α cell fractional area multiplied by the pancreatic weight, as previously described (34). In brief, seven sections per mouse, each separated by ∼25 μm, were immunostained for insulin and glucagon. For staining, sections were deparaffinized (2X Xylene for 5 minutes, 2X 100% Ethanol for 2 minutes, 95% Ethanol, 90% Ethanol, 80% Ethanol, and 70% Ethanol for 1 minute each, and 2X Milli-Q water for 2 minutes each), and endogenous peroxidase activity was quenched with 0.45% H_2_O_2_ (H325, Fisher Scientific) in 200 mL of methanol for 30 minutes. Reservoirs were drawn around tissue sections with an ImmEdge® Hydrophobic Barrier PAP Pen (H-4000, Vector Labs). Sections were rinsed in water, blocked with 2.5% horse blocking serum (MP-7401A, Vector Labs) for 30 minutes, and then incubated with primary anti-insulin antibody (3014S, Cell Signaling Technology) or primary anti-glucagon antibody (ab92517, Abcam) diluted 1:400 in 2.5% horse blocking serum overnight at 4 °C. Slides were washed in 1X PBS, then incubated with peroxidase-conjugated anti-rabbit Ig (MP-7401, Vector Laboratories) for 30 minutes. Slides were washed in 1X PBS and incubated with DAB Substrate Kit – Peroxidase HRP (SK-4100, Vector Laboratories) for approximately 2 minutes before quenching with diH2O. Lastly, sections were stained with Hematoxylin (1051740500, Sigma-Aldrich) for 15 sec and dehydrated (2X 70% Ethanol, 2X 95% Ethanol, 2X 100% Ethanol, and 2X Xylene for 10 sec each). Slides were mounted with Permount mounting media (SP15-100, Fisher) and scanned using an AxioScan Z1 scanning microscope (Carl Zeiss, RRID:SCR_002677). The fractional area of β and α cells was measured using ZEN 2 software (Carl Zeiss, RRID:SCR_013672).

### 2.8 mRNA Sequencing, Library Generation, and Data Analysis

Islets were isolated from CTR (n=4) and OVX (n=3) female mice following 5 weeks of HFD feeding (6 weeks post-surgery), and bulk RNA sequencing analysis was performed. After 24 hours of recovery in fresh media following islet isolation, total RNA was extracted from >100 hand-picked islets using the phenol-chloroform method (35). Total RNA samples were first evaluated for quantity and quality using Agilent Bioanalyzer 2100. The RNA integrity number (RIN) was 8.15±0.12 (mean±SD). One hundred nanograms of total RNA was used for library preparation with the KAPA mRNA Hyperprep Kit (KK8581, Roche). Each resulting uniquely dual-indexed library was quantified and quality assessed by Qubit and Agilent TapeStation. Multiple libraries were pooled in equal molarity. The pooled libraries were sequenced with 2×100bp paired-end configuration on an Illumina NovaSeq 6000 sequencer.

The sequencing data were first assessed using FastQC for quality control. All sequenced libraries were then mapped to the mouse genome (UCSC mm10) using STAR RNA-seq aligner. The reads distribution across the genome was assessed using bamutils (from ngsutils). Uniquely mapped sequencing reads were assigned to mm10 refGene genes using featureCounts. Differential expression (DE) analyses were performed using edgeR implemented in the Bioconductor package to identify differentially expressed mRNAs between CTR and OVX samples. Biological coefficients of variation between the samples were estimated using an empirical Bayes approach under the assumption that the data follows a negative binomial distribution. Low expression transcripts were filtered out based on percentage of samples (less than 50%) and CPM cutoff of 1. Statistical significance was defined as p-value <0.05 and a fold change (FC) >2 of expression level between comparison of CTR and OVX mice. The locus-by-locus volcano plots were generated using R package. DAVID was utilized for pathway enriched analysis for statistically significantly differentially expressed genes based on GO BP terms.

### 2.9 Quantification and Statistical Analysis

Statistical analysis was performed using Prism X software (GraphPad, RRID:SCR_002798). To compare two datasets, a two-tailed Student’s t-test was used. For experiments using 3 independent groups, data were analyzed by a one-way ANOVA. GTTs and ITTs were analyzed by two-way ANOVA followed by Tukey multiple comparison test. Results are reported as mean ± SD. P-value of less than 0.05 was considered to be a significant difference between groups.

## 3. Results

### 3.1 Age-related modulation in pathways related to β cell function in islets of female but not male human organ donors

To gain an unbiased understanding of the sex-specific pathways activated in the aging β cell, we queried a publicly available dataset (NCBI GEO dataset *GSE152111*), which included RNA sequencing results from 26 islet samples from female organ donors and 40 islet samples from male organ donors with information about the individual’s age, sex, and BMI. Because female samples did not have “menopausal status” listed as a variable, we instead chose to evaluate age as a continuous variable. We first determined genes that were significantly correlated with age in either sex. In this analysis, the expression of 1587 genes was significantly correlated with age in females (813 negatively and 774 positively, r > 0.4, p <0.05). In contrast, the expression of only 43 genes was correlated with age in males (18 negatively and 25 positively, r > 0.4, p <0.05). Of these genes, only five (*RBFOX1*, *CDKN2A*, *SPINK2*, *APMAP*, and *SULF2*) were common between the two sexes (**Fig. 1A**). Functional enrichment analysis of age-correlated genes revealed 312 pathways to be significantly enriched in the female islets, while only 65 pathways were significantly enriched in the males (fold enrichment ≥ 1.3 and p <0.05). Four pathways were shared between the sexes, including “skeletal muscle tissue development,” “striated muscle tissue development,” “steroid metabolic process,” and “skeletal muscle organ development” (**Fig. 1B**). Whereas β cell-specific pathways were largely absent from the male islets (**Fig. 1C**), in females, several pathways relevant to β cell function were identified, including “autophagy,” “response to insulin,” and “glucose transport” (**Fig. 1D**). Results from this analysis suggest a more pronounced effect of aging on gene expression in female compared to male islets and highlight age-related changes in genes associated with key β cell functions in female islets.

**Figure 1.**
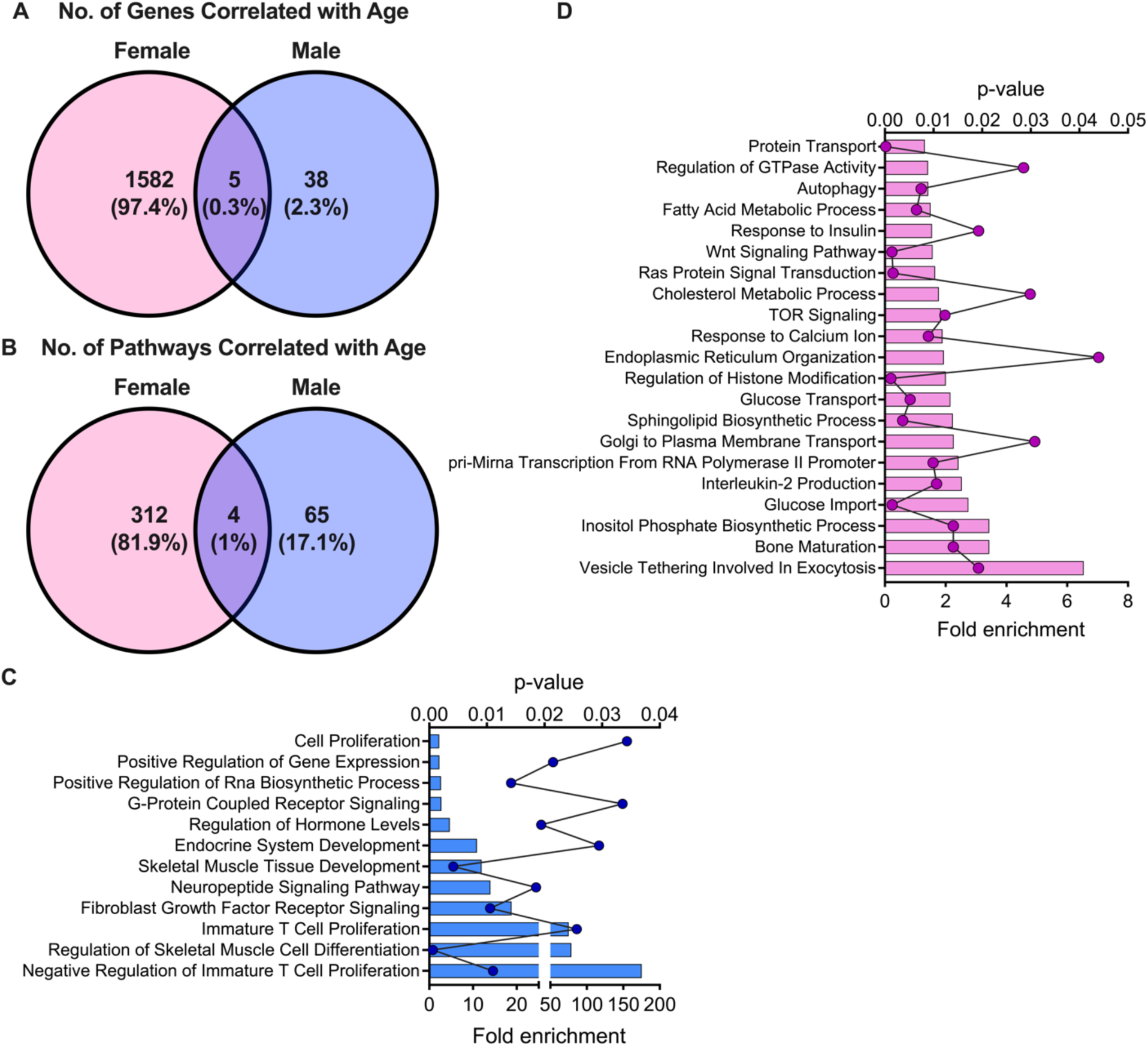
Correlation analysis using RNA-seq data from islet samples from human donors revealed significant age-related modulation in pathways related to β cell function in females but not males. NCBI GEO dataset (*GSE152111*), which consisted of islet RNA-seq results from female and male organ donors, was retrieved to assess the relationship between age and gene expression in males and females. A: Correlation analysis showing genes correlated with age in females (pink), males (blue), and both females and males (purple overlap). B-D: Functional enrichment analysis was performed using DAVID on the significantly correlated genes in the sexes and compared. The quantitative comparison is shown in (B) and representative pathways are illustrated for males (C) and females (D).

### 3.2 Ovariectomy increases weight gain, fat mass, lean mass, and total mass in HFD-fed female mice

We hypothesized that this significant age-related modulation of gene expression in female islets in humans was related to loss of E2 signaling. To model loss of E2 signaling experimentally and determine the impact of cessation of E2 signaling on β cell health, bilateral ovariectomy (OVX) was performed in wildtype (WT) female mice at 10 weeks of age. WT female mice of the same age that underwent sham surgery (CTR) were used as controls (schematic shown in **Fig. 2A**). Prior to surgery, no differences were seen between groups in either lean mass, fat mass, or total mass (**Fig. S1A-C**). Surgery effectiveness was confirmed 5 weeks post-surgery via liquid chromatography-mass spectroscopy, which showed significantly reduced serum E2 levels in female OVX mice compared to female CTR mice (**Fig. 2B**). One-week post-surgery, CTR and OVX mice were initiated on HFD containing 45% of calories from fat. After 4 weeks of HFD, female OVX mice exhibited increased lean mass (**Fig. 2C**), fat mass (**Fig. 2D**), total mass (**Fig. 2E**), and overall weight gain compared to female CTR mice (**Fig. 2F-G**).

**Figure 2.**
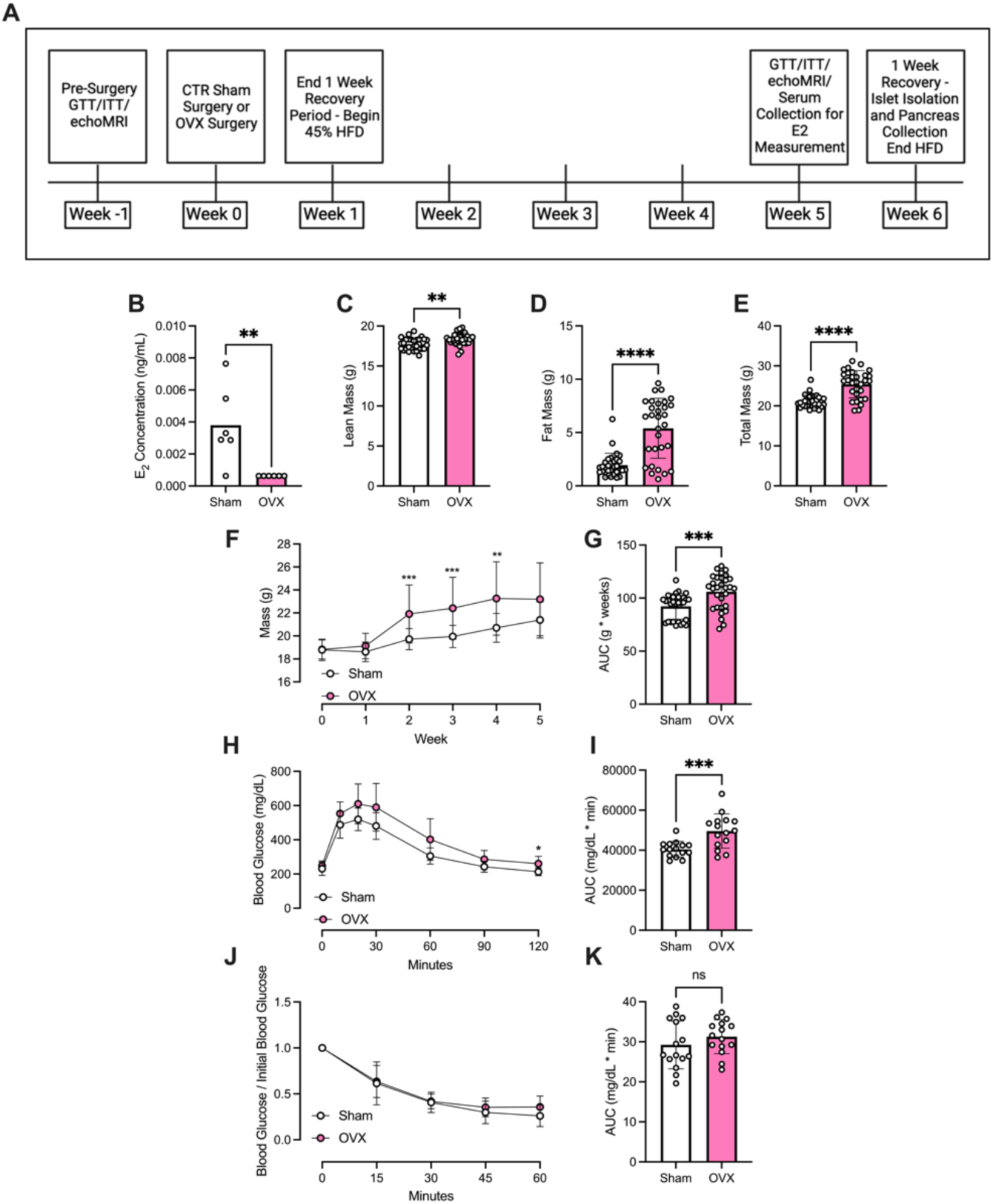
OVX in female mice causes weight gain, glucose intolerance, and insulin resistance upon HFD feeding. Control sham surgery (CTR) or ovariectomy surgery (OVX) was performed in WT C57Bl6 mice at 10 weeks of age. Control and OVX female mice were fed HFD for 4 weeks after 1 week of recovery from surgery. A: Experimental timeline. B-K: B: LC-MS where E2 concentrations less than the lower limit of detection (LOD) were recorded as half the lower LOD (2). Lean (C), fat (D), and total mass (E) were measured using the EchoMRI 500 Body Composition Analyzer. F-G: Changes in body weight were monitored over the HFD feeding period (F). AUC is shown graphically (G). H-I: Glucose tolerance was measured using a GTT (1.5 g/kg glucose dosed to lean mass) in HFD-fed CTR and OVX female mice (H). GTT results were analyzed with AUC analysis (I). ITT was performed (0.75 g/kg dosed to lean mass) (J). ITT results were analyzed using AUC (K). Replicates are indicated with circles; n ≥ 5 in each group. Results are displayed as mean ± SD. A two-tailed Student *t* test was used to compare the means between two groups. Weight gain, GTT, and ITT were analyzed with a two-way ANOVA followed by Tukey multiple comparison test. Indicated differences are statistically significant: \**P* <0.05, \*\**P* <0.01, \*\*\**P* <0.001, \*\*\*\**P* <0.0001.

### 3.3 Ovariectomy leads to glucose and insulin intolerance in female HFD-fed mice

Prior to surgery, IP GTTs and ITTs showed no difference between groups in systemic glucose tolerance or peripheral insulin sensitivity (**Fig. S1D-G**). However, after 4 weeks of HFD, female HFD-fed OVX mice had increased glucose excursions and worsened glucose tolerance when compared to female HFD-fed CTR mice (**Fig. 2H-I**). Notably, at this time point, insulin tolerance was not different between the groups, suggesting early modulation of β cell function without measurable differences in insulin sensitivity (**Fig. 2J-K**).

### 3.4 RNA-seq analysis reveals significant modulation in pathways related to β cell function in islets from OVX-female mice

To obtain mechanistic insights into the pathways leading to glucose intolerance in the OVX model, islets were isolated from HFD-fed female CTR and OVX mice and subjected to RNA sequencing analysis. Principal component analysis revealed a distinct separation between CTR and OVX mice (**Fig. 3A**). We identified 292 differentially expressed genes based on a threshold fold-change ≥2 and p-value <0.05. Of these differentially expressed genes, 185 genes were downregulated and 107 genes were upregulated. Expression patterns of the top 50 differentially expressed genes are shown in **Fig. 3B**. Pathway analysis identified 76 pathways that were significantly modulated (*p* <0.05), including several pathways relevant to β cell function including “glucose homeostasis,” “type B pancreatic cell proliferation,” and “endocrine pancreas development” (**Fig. 3C**). Notably, within these pathways, several downregulated genes including *Nkx6.1, Pdx1,* and *Ucn3* were associated with the maintenance of β cell identity (**Fig. 3D**).

**Figure 3.**
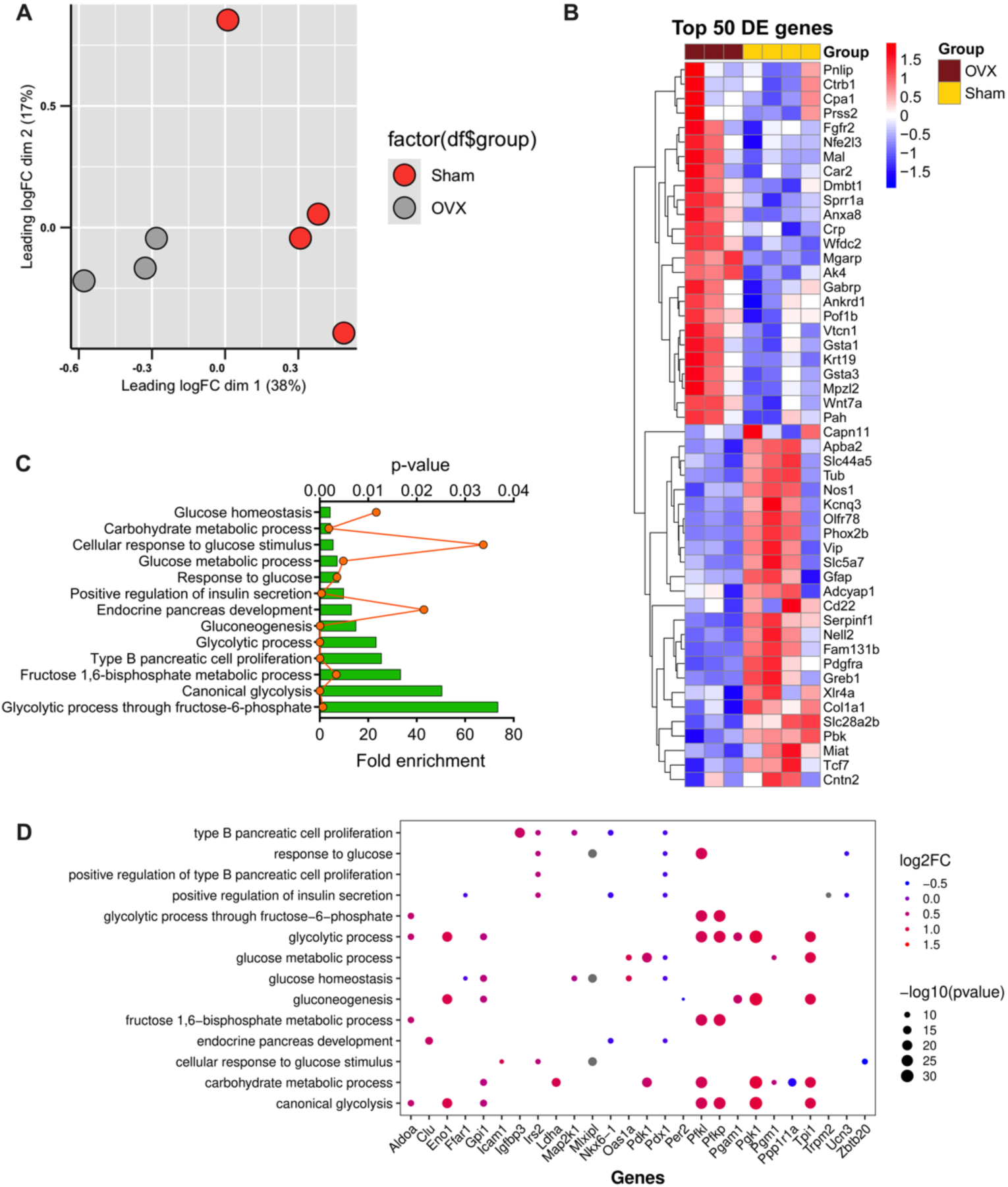
RNA-Seq analysis revealed differential expression of genes related to loss of β cell identity and function in islets from female OVX mice. Pancreatic islets from HFD-fed CTR and OVX female mice were subjected to RNA-sequencing analysis using 100 ng of total RNA per sample. A: Principal component analysis revealed clear separation between CTR and OVX islets. Log cpm values were used to generate the PCA plot. B: Top 50 differentially expressed genes are shown in a heat map. Genes in blue indicate downregulated genes in OVX islets and genes in red indicate upregulated genes in OVX islets. C: Functional enrichment analysis was performed using DAVID; 13 pathways relevant to β cell function are illustrated. D: Dot plot illustrating the gene expression patterns within the pathways depicted in (C). A blue circle indicates a negative log fold change of the gene while a red circle indicates a larger, positive log fold change. Additionally, the size of the circle represents the -log10(p-value).

### 3.5 Ovariectomy increases α cell mass and decreases β cell identity gene expression in HFD-fed female mice

Our RNA-sequencing profile suggested that loss of E2 signaling via OVX may impair β cell function in female mice, potentially due to loss of β cell maturity. To test whether these changes were associated with alterations in endocrine cell composition, β and α cell mass were quantified in pancreatic sections from female CTR and OVX mice after 5 weeks of HFD. While there was not a significant change in β cell mass between groups, there was a significant increase in α cell mass in OVX compared to CTR mice (**Fig. 4A-B**). Consistent with this finding, immunofluorescence staining showed a significant increase in glucagon staining with no significant change in insulin staining (**Fig. 4C-E**). To assess whether changes in endocrine composition were related to changes in proliferation, we performed immunofluorescence staining to evaluate Ki67-positive nuclei present in insulin-positive cells or insulin-negative cells in both CTR and OVX mice (**Fig. 4F**). Analysis showed no significant change in Ki67-positive, insulin-positive nuclei (**Fig. 4G**) or Ki67-positive, insulin-negative nuclei (**Fig. 4H**). Given that there was no change in β cell mass, we next measured the expression of a panel of genes associated with β cell function and maturity in islets isolated from female HFD-fed CTR and OVX mice. We found a significant decrease in *Mafa*, *Ucn3*, and *Pdx1* gene expression in islets of HFD-fed OVX mice compared to control (**Fig. 4I**), consistent with changes observed in the RNA-sequencing results.

**Figure 4.**
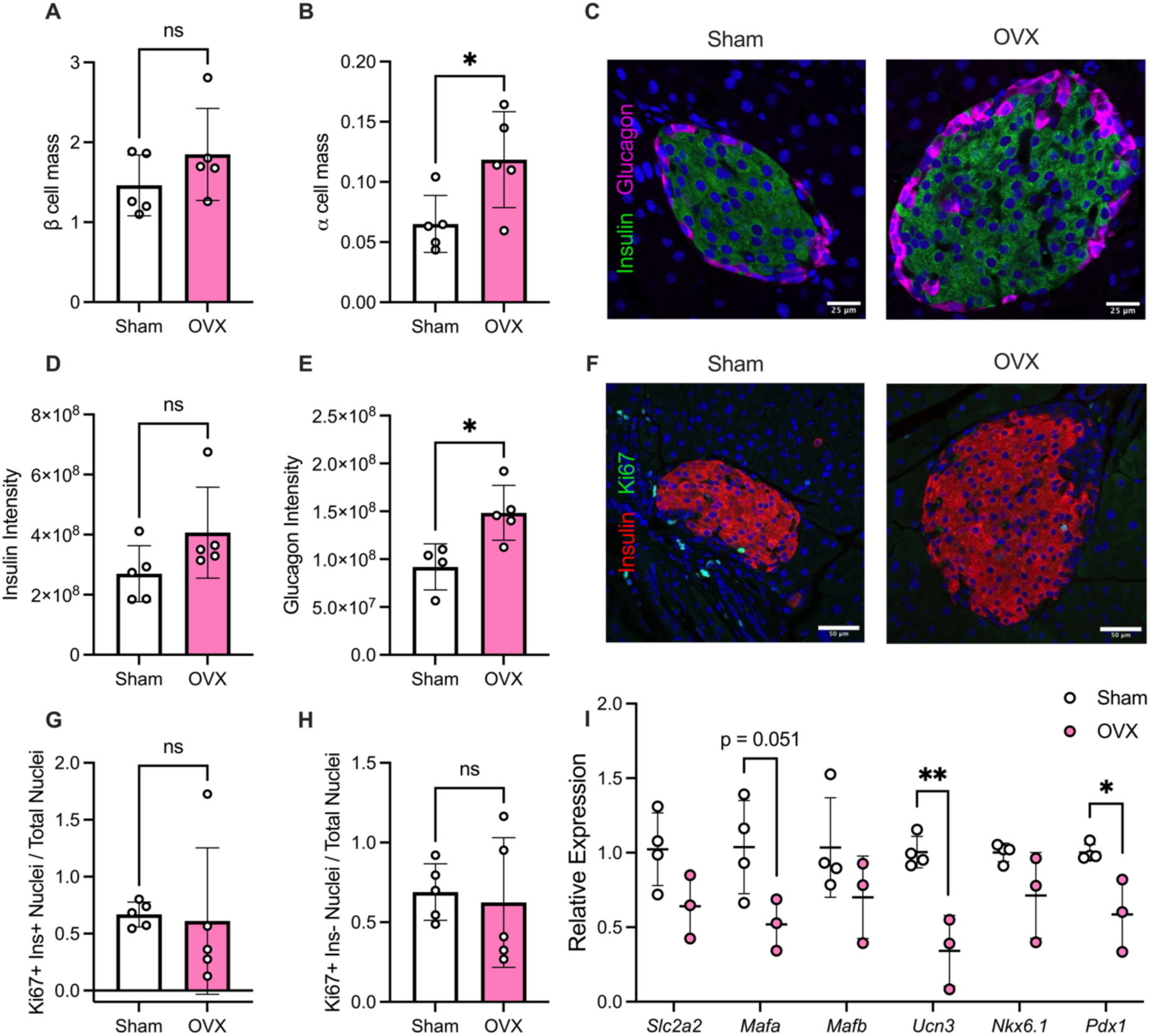
Loss of E2 signaling in female mice leads to loss of β cell identity and increased α cell mass under HFD conditions. A-B: The total mass of insulin-positive β cells and glucagon- positive α cells within pancreata was quantified in paraffin tissue sections by immunostaining for insulin and glucagon in female mice fed HFD for 5 weeks. C: Representative immunofluorescence images of islets in HFD-fed CTR and OVX female mice; sections were stained with anti-insulin (green) and anti-glucagon (magenta) antibodies. D-E: Integrated insulin (D) and glucagon (E) density was quantified in paraffin tissue sections by immunofluorescence staining for insulin and glucagon in female mice fed HFD for 5 weeks. F: Representative immunofluorescence images of islets in HFD-fed CTR and OVX female mice; sections were stained with anti-insulin (red) and anti-Ki67 (green) antibodies. G-H: Number of Ki67-positive nuclei were quantified in insulin-positive nuclei (G) and insulin-negative nuclei (H) and normalized to the total number of nuclei. I: Islets were isolated from HFD-fed CTR and OVX female mice. mRNA levels of *Slc2a2, Mafa, Mafb, Ucn3, Nkx6.1,* and *Pdx1* were analyzed by RT-qPCR. Results were normalized to *Actb* levels. Replicates are indicated with circles; n ≥ 3 in each group. Results are displayed as mean ± SD. A two-tailed Student *t* test was used to compare the means between two groups and a one-way ANOVA was used to compare the means between three or more independent groups followed by Tukey multiple comparison test. Indicated differences are statistically significant: \**P* <0.05, \*\**P* <0.01, \*\*\**P* <0.001, \*\*\*\**P* <0.0001.

### 3.6 Modulation of E2 signaling rescues loss of β cell identity gene expression in islets from OVX- female mice

E2 acts through three different receptors: ERα, ERβ, and GPER. We examined expression patterns of the three estrogen receptors in isolated islets, and we found that expression of *Esr1* and *Gper1* was significantly decreased in HFD-fed OVX mice compared to CTR mice while *Esr2* trends toward a decrease (**Fig. 5A-C**). Our current findings show that loss of E2 signaling via OVX decreases islet expression of genes associated with β cell function and maturity. Previously, we showed that β cell-specific loss of STIM1 led to reduced GPER expression and decreased β cell identity in female mice (25). Therefore, we next aimed to determine whether modulating E2 signaling could rescue the loss of β cell identity in an *ex vivo* setting. Islets isolated from female HFD-fed CTR and OVX mice were treated with E2, which activates ERα, ERβ, and GPER, or the GPER-specific agonist G-1 for 24 hours. G-1 significantly increased *Mafa*, *Ucn3*, and *Pdx1* gene expression in the OVX group, while E2 treatment significantly increased only *Ucn3* gene expression, suggesting that GPER may play a more specific role in the maintenance of β cell identity in female mice (**Fig. 5D**).

**Figure 5.**
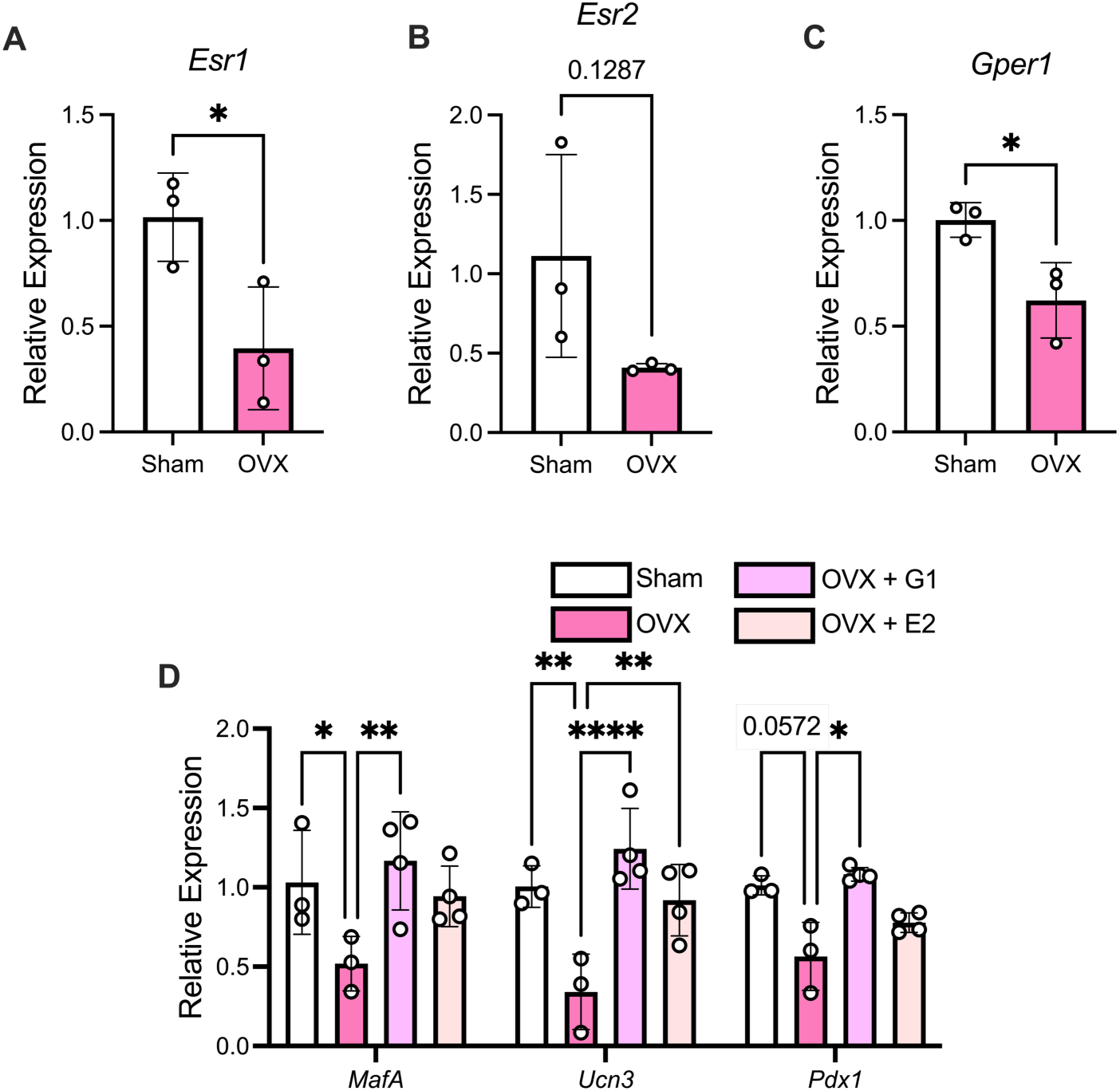
GPER agonism in OVX islets leads to a partial restoration of β cell identity gene expression. A-C: Islets were isolated from HFD-fed CTR and OVX female mice. mRNA levels of *Esr1* (A)*, Esr2* (B), and *Gper1* (C) were analyzed by RT-qPCR. Results were normalized to *Actb* levels. D: Islets isolated from HFD-fed CTR and OVX mice were treated *ex vivo* with G-1 or E2 for 24 hours, and mRNA levels of *Mafa, Ucn3*, and *Pdx1* were analyzed with RT-qPCR. Results were normalized to *Actb* levels. Replicates are indicated with circles; n ≥ 3 in each group. Results are displayed as mean ± SD. A two-way ANOVA was used to compare the groups followed by Tukey multiple comparison test. Indicated differences are statistically significant: \**P* <0.05, \*\**P* <0.01, \*\*\**P* <0.001, \*\*\*\**P* <0.0001.

## 4. Discussion

Epidemiological data highlight an important and time-dependent role for biological sex in the risk of T2D development across the lifespan. Consistent with this data, clinical and preclinical studies suggest that the specific milieu of sex hormones present in males and females influences aging-associated changes in metabolism (36). While the prevalence of T2D increases with age in both men and women, in young and middle-aged populations, men show a higher prevalence of T2D than age-matched females (1, 37). However, after undergoing menopause, women exhibit an increased risk for T2D that is comparable to the risk seen for age-matched men (10). When looking specifically at age-related differences in T2D development between the sexes, as women with impaired glucose metabolism age, it has been shown that their β cell function declines while aging males with T2D show deterioration of glucose metabolism (38). Brownrigg et al. expanded on these findings by demonstrating that a baseline sex difference in β cell gene expression likely underlies the sex-specific transcriptional responses to perturbations like ER stress and T2D (39). Menopause in females demarcates the period of reproductive aging that manifests as low ovarian hormone secretion (40). In the United States, menopause in women occurs at an average age of 51, marks a sudden sharp decline in estrogens and progesterone, and is a major driver of the female aging process. However, this drastic hormonal change does not occur in males, and the analogous process, called andropause, is more difficult to define due to testosterone levels decreasing at an average rate of only 1% per year (41). To evaluate how aging and loss of E2 signaling impacts the female islet, we analyzed a publicly available RNA-seq dataset of 40 males and 26 females, including 15 females greater than or equal to 51 years of age (the average age of menopause). We found significant differences between aging female and male islets. Remarkably, the expression of 1582 genes in females versus only 38 genes in males was correlated with advancing age in islets from human organ donors. Consistent with this finding, pathway analysis demonstrated that 312 pathways were significantly modulated in females, while only 65 pathways were modulated in males. We hypothesize that this sex-specific difference may be due to the loss of E2 signaling in females that causes a significant change in β cell function with age, while the more gradual decline in androgens in males may not have as large an impact. Notably, we found that many pathways correlated with age in female islets were relevant to β cell function (“autophagy,” response to insulin,” and “glucose transport”), while β cell-specific pathways were absent in males.

Interestingly, Shrestha et al. examined a published single-cell (sc) RNA-sequencing dataset of human pancreatic islets containing 68 donors without diabetes and 35 donors with T2D. They found that β cells from younger (0-29 years of age) or older (> 60 years of age) donors had higher expression of genes playing a role in immune response, response to ER stress, autophagy, and fatty acid metabolism when compared to β cells from middle-aged donors, corroborating the results we saw in aging female islets (42). However, this group did not account for biological sex as a variable in the aging process. In a preprint by Arrojo e Drigo et al., the investigators similarly completed a meta-analysis of three scRNA-seq datasets derived from human islet cells from nondiabetic and diabetic donors. They divided the sample cohort into three age ranges: 1) <6 years of age, 2) 20-48 years of age, and 3) 54-68 years of age and investigated sex differences in the aging β cell. Aged male β cells exhibited upregulation of genes associated with other endocrine cell types, but these changes were not seen in aged female β cells which expressed high levels of genes involved in β cell function, identity, and glucose metabolism, suggesting that cell function is maintained (9). However, this study only included one female donor greater than age 50, making it difficult to thoroughly examine the effect of menopause. In contrast, our dataset contained 15 female donors greater than or equal to 51 years of age, with 26 total female donors. This provided us the opportunity to gain insights into how the female β cell changes over a span of ages (16 to 61 years of age) and evaluate how E2 may play a role in this process.

Based on the differences we observed in the aging female and male human islets, we next evaluated the specific role of E2 in β cell health by removing E2 signaling in an OVX mouse model. Previous studies have shown that OVX in mice impairs glucose tolerance and causes insulin resistance. Camporez et al. performed OVX surgery in female mice at 8 weeks of age followed by 4 weeks of HFD feeding in parallel with vehicle or E2 injection. OVX mice displayed reduced whole-body energy expenditure, impaired glucose tolerance, and whole-body insulin resistance. Importantly, hepatic and muscle insulin resistance were improved when OVX mice were treated with E2 injections (43). Fewer studies have examined the impact of OVX on β cell function. De Paoli et al. performed OVX surgeries in an Akita mouse model of hyperglycemia-induced atherosclerosis to analyze the effects of E2 on the maintenance of β cell health and function and atherosclerosis progression. They found that OVX mice developed chronic hyperglycemia and reduced β cell mass, while exogenous E2 supplementation via slow-release E2 pellets for 90 days rescued hyperglycemia by normalizing blood glucose levels and maintaining β cell mass (44).

Our data corroborates many of these findings and adds important information about how OVX impacts β cell health in a model of diet-induced obesity. In our study, we also observed that OVX impaired glucose tolerance. In contrast to the Camporez study, we did not detect a significant different in insulin sensitivity, and we did not see a change in β cell mass as observed in the De Paoli study. Interestingly, we observed increased α cell mass with no change in α or β cell proliferation. Although there was no change in β cell mass in our HFD OVX model, there was a significant decrease in the expression of key β cell identity genes (*Mafa*, *Ucn3*, and *Pdx1*) in OVX mice compared to CTR mice. In the analysis of the human pancreatic islets scRNA-seq dataset completed by Shrestha et al. (described above), they also found that β cells from younger or older donors had lower expression levels of β cell identity gene markers, in agreement with our findings (45). Similarly, previous studies have shown reduced protein expression of these markers of β cell identity and maturity in T2D (46); however, this study did not report the sex of donor islets. Interestingly, other studies have implicated de-differentiation as a causative molecular pathway in T2D pathogenesis and this phenomenon was described in aged female mice (47, 48).

When we treated isolated islets from OVX-HFD mice *ex vivo* with E2 or G-1, there was a significant rescue of β cell identity gene expression. Several studies have linked GPER signaling to β cell function. Mårtensson et al. showed that female whole-body GPER KO mice had hyperglycemia and impaired glucose tolerance (49). GPER KO has been shown also to increase the susceptibility of female mice to streptozotocin (STZ)-induced diabetes (50), and Jacovetti et al. found that activation of GPER via G-1 decreases miR-338-3p expression and results in an expansion of β cell mass and increased β cell proliferation (51). Lastly, Lombardi et al. showed that pretreatment of islets with E2 rescued glucotoxicity-induced β cell dedifferentiation, and co-treatment with E2 and the GPER antagonist G15 reversed this effect (52). These previous findings combined with our data indicate that E2 signaling through GPER may maintain β cell health through the conservation of β cell identity.

There are some limitations to our study that should be acknowledged. First, for the human sample data, menopausal status of the female samples was not readily available. Because women undergo menopause at variable ages, this correlation analysis may not entirely exemplify how loss of E2 signaling correlates with β cell identity gene expression. Second, findings in our female OVX HFD model may be confounded by effects of obesity on peripheral insulin sensitivity and inflammation. While we did not observed differences by ITT, it is possible a more sensitive measure of insulin resistance would detect differences at this early timepoint.

Notwithstanding these limitations, we have shown that upon abrupt loss of endogenous E2 signaling via OVX surgery, HFD-fed female mice exhibit increased glucose excursions and overall weight gain compared to HFD-fed female CTR mice. HFD-fed OVX mice had an increase in α cell mass and a decrease in β cell identity gene expression, and treatment of OVX-islets *ex vivo* with the GPER agonist G-1 rescued the loss of β cell identity gene expression. Taken together, these data suggest that E2 signaling through GPER plays a critical role in the maintenance of β cell health and identity and the pathogenesis of T2D in females.

## Acknowledgements

We thank the Wisconsin National Primate Research Center Assay Services Unit (University of Wisconsin) for performing liquid chromatography-mass spectroscopy and Dr. Emily Anderson-Baucum (Indiana University School of Medicine) for her helpful edits.

## CRediT Authorship Contribution Statement

MRM, TK, KLK, and CEM conceived and designed the study. MRM, CR, KO, and LU performed experiments. AF performed the OVX surgeries. JDCA performed the β and α cell mass analysis. PK and WW completed the human islet sequencing analysis and the RNA-seq analysis. MRM performed the statistical analysis. MRM and CEM interpreted the data. MRM and CEM wrote the manuscript, and PK and WW assisted with the writing of the methods. All authors provided critical revisions and edits to the manuscript. All authors read and approved of the final manuscript. CEM is the guarantor of this work. Both MRM and CEM have verified the underlying data of this manuscript.

## Data Availability

Data presented in this manuscript are available from the corresponding author upon request.

