## Supplemental Figures for "G Protein Coupled Estrogen Receptor Signaling Maintains β Cell Identity in Female Mice"

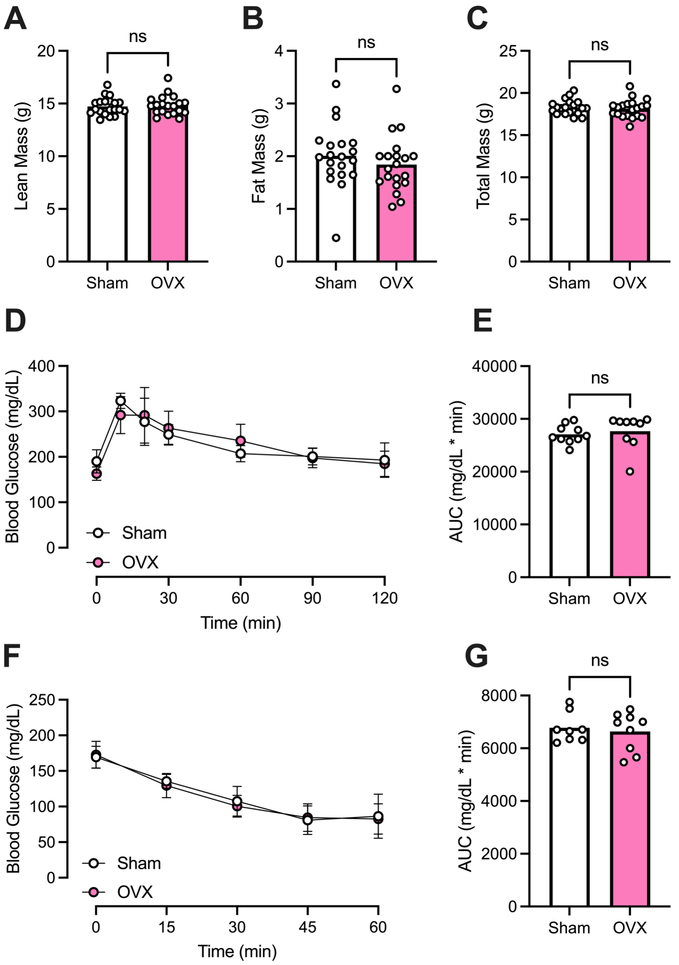


**Figure S1. No differences in body composition, glucose tolerance, or insulin tolerance between female mice prior to CTR or OVX surgery.** At 9 weeks of age, prior to control sham (CTR) or ovariectomy (OVX) surgery, body composition, glucose tolerance, and insulin tolerance were evaluated in WT C57Bl6 female mice. A-C: Lean (A), fat (B), and total mass (C) were measured using the EchoMRI 500 Body Composition Analyzer. D-E: Glucose tolerance was measured using a GTT (1.5 g/kg glucose dosed to lean mass) (D). GTT results were analyzed with AUC analysis (E). ITT was performed (0.75 g/kg dosed to lean mass) (F). ITT results were analyzed using AUC (G). Replicates are indicated with circles; $n \geq 5$ in each group. Results are displayed as mean ± SD. A two-tailed Student *t* test was used to compare the means between two groups. Weight gain, GTT, and ITT were analyzed with a two-way ANOVA followed by Tukey multiple comparison test.
